# Molecular basis for dual functions in pilus assembly modulated by the lid of a pilus-specific sortase

**DOI:** 10.1101/2023.11.05.565703

**Authors:** Chungyu Chang, HyLam Ton-That, Jerzy Osipiuk, Andrzej Joachimiak, Asis Das, Hung Ton-That

**Author notes:** For correspondence: Hung Ton-That.

## Abstract

The biphasic assembly of Gram-positive pili begins with the covalent polymerization of distinct pilins catalyzed by a pilus-specific sortase, followed by the cell wall anchoring of the resulting polymers mediated by the housekeeping sortase. In *Actinomyces oris*, the pilus-specific sortase SrtC2 not only polymerizes FimA pilins to assemble type 2 fimbriae with CafA at the tip, but it can also act as the anchoring sortase, linking both FimA polymers and SrtC1-catalyzed FimP polymers (type 1 fimbriae) to peptidoglycan when the housekeeping sortase SrtA is inactive. To date, the structure-function determinants governing the unique substrate specificity and dual enzymatic activity of SrtC2 have not been illuminated. Here, we present the crystal structure of SrtC2 solved to 2.10-Å resolution. SrtC2 harbors a canonical sortase fold and a lid typical for class C sortases and additional features specific to SrtC2. Structural, biochemical, and mutational analyses of SrtC2 reveal that the extended lid of SrtC2 modulates its dual activity. Specifically, we demonstrate that the polymerizing activity of SrtC2 is still maintained by alanine-substitution, partial deletion, and replacement of the SrtC2 lid with the SrtC1 lid. Strikingly, pilus incorporation of CafA is significantly reduced by these mutations, leading to compromised polymicrobial interactions mediated by CafA. In a *srtA* mutant, the partial deletion of the SrtC2 lid reduces surface anchoring of FimP polymers, and the lid-swapping mutation enhances this process, while both mutations diminish surface anchoring of FimA pili. Evidently, the extended lid of SrtC2 enables the enzyme the cell wall-anchoring activity in a substrate-selective fashion.

## INTRODUCTION

Covalently crosslinked pilus polymers (also known as pili or fimbriae) are found on the surface of many Gram-positive bacteria, including various pathogens such as *Corynebacterium diphtheriae*, *Actinomyces oris*, *Bacillus cereus*, *Enterococcus faecalis* and many species of streptococci (1–7). The model organism in the present study is *A. oris*, an actinobacterium residing in the human oral cavity which utilizes two antigenically distinct types of fimbriae that perform unique functions in colonization, biofilm formation, and polymicrobial interactions known as coaggregation. The type 1 fimbriae, comprised of the fimbrial shaft protein FimP and the tip protein FimQ (8), mediate bacterial adherence to proline-rich proteins that coat the tooth surface (9,10), while the type 2 fimbriae, made of the fimbrial shaft FimA and the tip fimbrillin FimB (8,11), promote biofilm formation and coaggregation (11–13). Importantly, there are two distinct forms of the heterodimeric type 2 fimbriae; both are made of the same shaft fimbrillin FimA, but their tip fimbrillin is distinct – one is FimB and the other CafA. Functionally, FimA mediates biofilm formation while CafA is essential for coaggregation (14); however, the function of FimB is unknown.

*A. oris* shares with other Gram-positive bacteria a conserved biphasic mechanism of sortase-catalyzed pilus assembly first described in *C. diphtheriae* (1). It consists of the pilus polymerization step in the first phase involving the covalent linking of pilin monomers catalyzed by a pilus-specific sortase (class C), followed by the anchoring of resulting pilus polymers to the cell wall by the ‘housekeeping’ sortase (class A or E) that (as its name implies) acts to attach many distinct antigens to the bacterial cell wall for surface display (15). Precursors of these pilins harbor a C-terminal cell wall sorting signal (CWSS), comprised of a LPXTG motif preceding a hydrophobic domain and a positively charged tail (16). With the type 2 fimbriae as an example, a model of pilus assembly has been proposed (15); in this biphasic model, the pilus-specific sortase SrtC2 catalyzes pilus polymerization, first linking the tip pilin CafA to a FimA pilin and then connecting this dimer to individual FimA pilins sequentially via repetitive transpeptidation reactions. In these transpeptidation reactions, SrtC2 cleaves the LPXTG motif between threonine and glycine and links the terminal threonine residue to the nucleophilic lysine residue within the pilin motif of FimA. The housekeeping sortase SrtA then captures the resulting polymers and links them to the peptidoglycan, likely via the stem peptide of lipid II precursors and/or the uncrosslinked cell wall as proposed for *E*. *faecalis* pili (17). The pilus polymerization activity of SrtC2 appears to be highly specific for type 2 fimbriae. That is, SrtC2 cannot substitute the pilus-specific sortase SrtC1 for polymerization of type 1 fimbriae, and vice versa (8,18). On the other hand, the housekeeping sortase SrtA functions indiscriminately on FimA and FimP polymers, anchoring both to the bacterial cell wall, in addition to many other surface proteins that harbor a conserved LPXTG (18). In this context, it is noteworthy that both FimA and FimP possess the LPLTG motif, while many other surface proteins contain the LAXTG motif (8,14).

To date, the critical questions of what structural determinants govern the exquisite substrate specificity of the pilus-specific sortases SrtC1 and SrtC2 and what enables the housekeeping sortase SrtA to anchor a diverse set of substrates to the cell wall have not been addressed. Previous X-ray crystallographic determination of the structures of SrtA and SrtC1 revealed several conserved features of the sortase family, both having an 8-stranded β-barrel, now known as the sortase fold, with β7 and β8 containing the catalytic site comprised of a His-Cys-Arg triad (18,19). Intriguingly, SrtC1 harbors a so-called structural lid that covers the catalytic site and is absent in all housekeeping sortases including *A. oris* SrtA (20,21). This lid has been postulated to play a role in conferring substrate specificity to sortase enzymes (21,22). The presence of a similar lid in SrtC2 has been revealed by structural modeling (13). However, while alanine-substitution of the conserved His and Cys residues in SrtC2 abrogates pilus polymerization of type 2 fimbriae, mutations of two residues within the predicted lid region of SrtC2 do not affect assembly of type 2 fimbriae (13). It is noteworthy that *srtA* is an essential gene in *A. oris* as genetic disruption of *srtA* causes excessive membrane accumulation of the glycoprotein GspA, leading to cell death (23). In a viable mutant strain devoid of the essential gene *srtA* and *gspA* (a genetic suppressor of *srtA* deletion), a significant portion of both FimA and FimP pilus polymers are still anchored to the bacterial cell wall, while a substantial fraction of both pili fails to be linked to the cell wall and consequently are released into the extracellular milieu, as expected from a pivotal role of the housekeeping sortase in pilus anchoring (23). These results led to the inescapable conclusion that in the absence of SrtA, either SrtC1 or SrtC2 catalyzes the anchoring of pilus polymers to peptidoglycan, albeit with poor efficiency compared to SrtA. Subsequent studies revealed that it is SrtC2, and not SrtC1, which is required for this cell wall anchoring function in the absence of SrtA (18). To date, however, what determines the dual activities of a sortase such as SrtC2, i.e. both pilus polymerization and cell wall anchoring, has remained a mystery.

Here, we present an X-ray crystal structure of SrtC2 solved to 2.10-Å resolution, which unveiled the conserved and distinct features of SrtC2, relative to SrtC1 and SrtA. By a combination of structural comparison, biochemical studies, domain swapping and mutational analyses, we demonstrate for the first time that the lid of SrtC2 itself modulates its dual enzymatic activity, a conclusion that represents a significant advance to our understanding of the structural basis of sortase-mediated surface assembly in Gram-positive bacteria.

## RESULTS

### Structural determination of the *A. oris* pilus-specific sortase SrtC2

In a previous genetic study (18), we discovered that *A. oris* SrtC2 possesses both pilus polymerization and cell wall anchoring functions, apparent when the housekeeping sortase SrtA is absent. SrtC2 not only anchors the cognate FimA pilus polymers, which it polymerizes, but also the FimP polymers that are assembled by the pilus-specific sortase SrtC1 (10,18). To gain insights into the molecular basis for this dual activity of SrtC2, we attempted to determine its structure by X-ray crystallography. The best needle-like crystals of the SrtC2 protein diffracted up to 2.10-Å resolution in space group P6_2_22. The final structure shows an excellent refinement statistics with R-work and R-free factors of 16.7 and 20.1 %, respectively (Table 1). The structure is a continuous amino-acid chain that spans residues 82-267, with the N-terminal residues 60-81 and C-terminal residues 268-282 found to be disordered. In the crystal, the protein is a monomer without any indication of forming a protein dimer as it was previously reported for *A. oris* SrtC1 (19).

The SrtC2 structure harbors a canonical sortase fold (24–27), with a core consisting of eight-strand β-barrel flanked by two α-and three 3_10_-helices (Fig. 1A and Fig. 2). The active site is located at the top of the β-barrel, containing the conserved catalytic triad H^184^-C^246^-R^255^. Similar to other pilus-specific sortases, the active site of SrtC2 is partially covered by a lid (residues 97-122) that seemingly occludes access to catalytic residues C^246^ and R^255^. This 26-residue lid is stabilized in the structure by multiple hydrogen bonds between the lid loop (residues 104-108) and the core loop (residues 246-255), including bonding between the active site R255 and main-chain oxygen of E^104^ lid residue (Fig. 1B). Additional lid bonding is created by a small β-sheet formed by β11 core strand (residues 230-233) and two short β2 and β3 lid strands (residues 100-103 and 111-112, respectively) (Fig. 1B). Interestingly, the presence of β11 strand appears to be unique for *A. oris* SrtC2, as α-helices, in place of this strand, are found in many other sortases, with the exception of a twin non-canonical ‘open-form’ in *Streptococcus agalactiae* SrtC1, having a β-turn in this location (Fig. 2). The lid conformation of *A. oris* SrtC2 is also distinctive. The β2-β3 strand interaction and the presence of small β1-β4 sheet (residues 97-98 and 115-116, respectively) seem to be responsible for a significantly slimmer conformation compared to the majority of sortases, including *A. oris* SrtC1 (Fig. 1B-1C, Fig. 2, and Fig. 3A).

**Figure 1:**
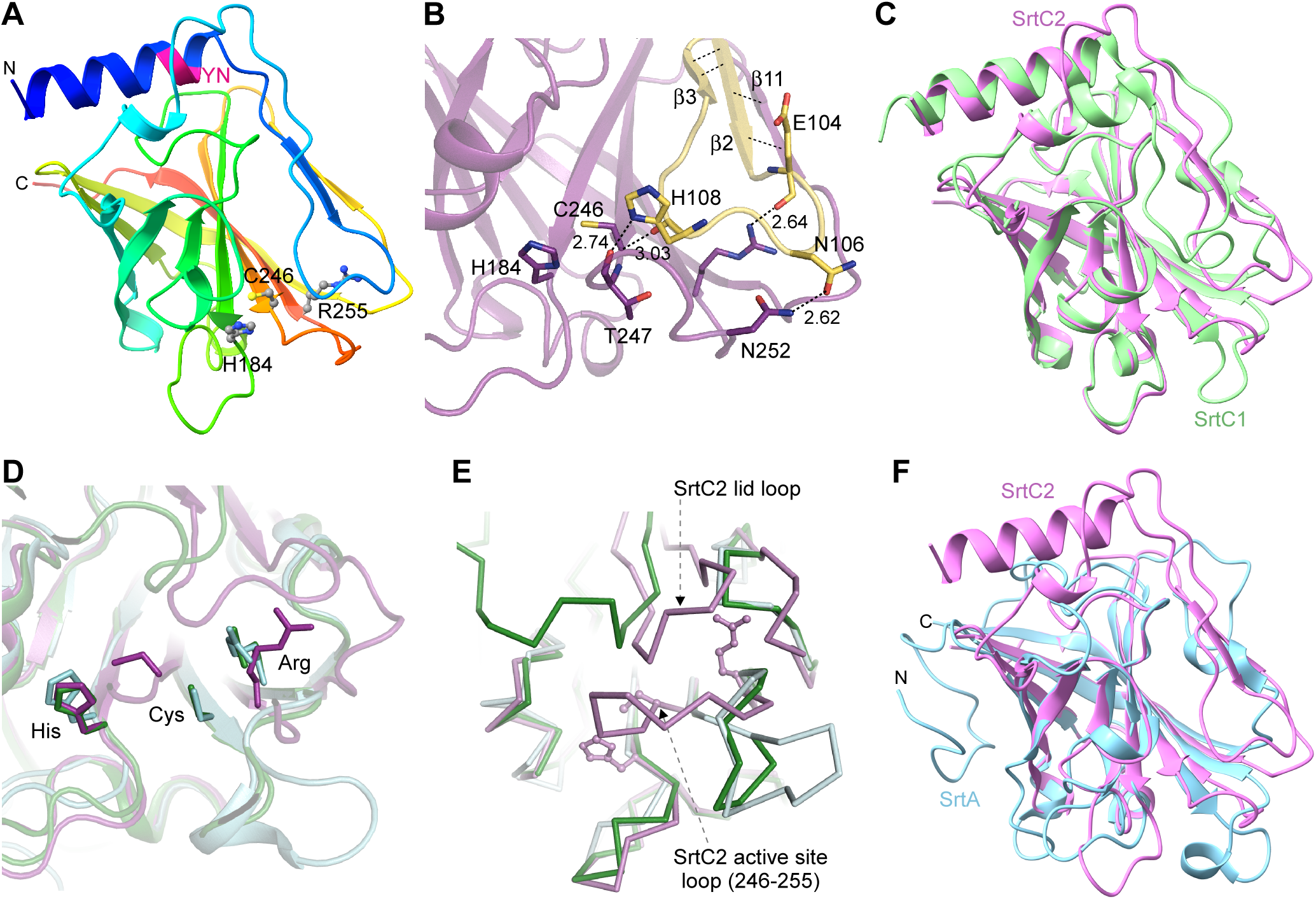
Crystal structural analysis of *A. oris* sortase SrtC2. **(A)** The crystal structure of *A. oris* pilin-specific sortase C2 (residue 82 to 267) was determined to 2.14 Å resolution. Presented is a rainbow cartoon, with a flexible lid (light blue) covering the three catalytic residues H184, C246, and R255 shown in ball-and-sticks. **(B)** Enlarged is the catalytic pocket of SrtC2 with key interactions between the lid (yellow) and the protein core (pink). **(C)** The SrtC2 structure (pink) is superimposed with the *A. oris* SrtC1 structure (2XWG) (green). **(D)** Presented is superposition of active centers of the *A. oris* SrtC2 structure (pink), *A. oris* SrtC1 (green) and *A. oris* SrtA (5UTT; blue). **(E)** The active center and lid region of SrtC2 (pink) is superimposed with that of A. oris SrtC1 (green). (F) The SrtC2 structure (pink) is superimposed with the *A. oris* SrtA structure (5UTT) (blue).

**Figure 2:**
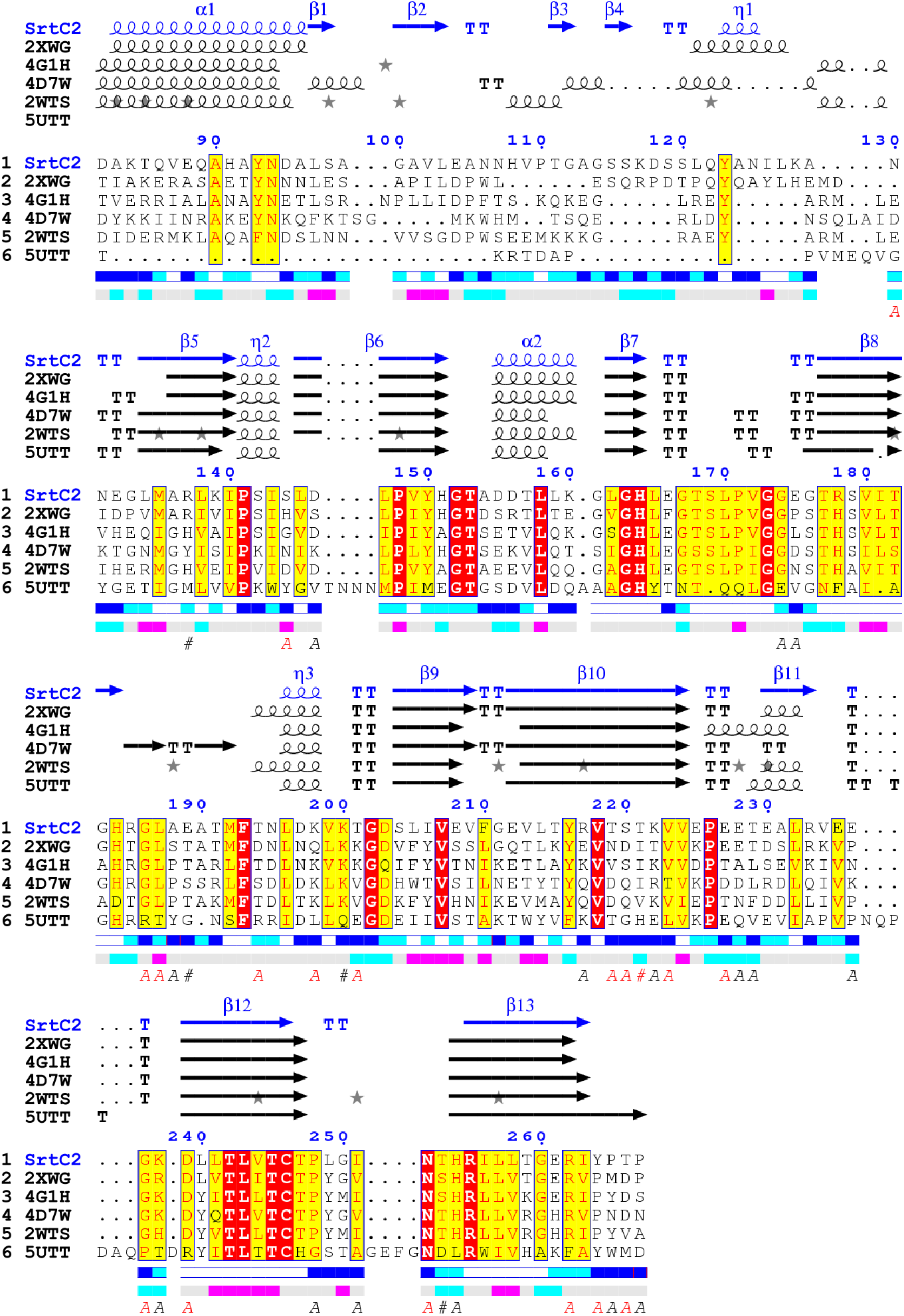
Sequence alignments of *A. oris* SrtC2 structural homologs. Sequence alignments were performed with ENDscript (45) for the following structures (top to bottom): *A. oris* SrtC2, *A. oris* SrtC1 (2XWG), *S. agalactiae* SrtC2 (4G1H), *S. agalactiae* SrtC1 (4D7W), *S. pneumoniae* SrtC1 (2WTS) and *A. oris* SrtA (5UTT). The protein secondary structures are shown above the alignments as α# for α-helices, β#-β-strands, ƞ – 3_10_ helices, TT – β-turns. Solvent accessibility is rendered by a first bar below the sequence (blue is accessible, cyan is intermediate, white is buried) and hydropathy by a second bar below (pink is hydrophobic, white is neutral, cyan is hydrophilic).

**Figure 3:**
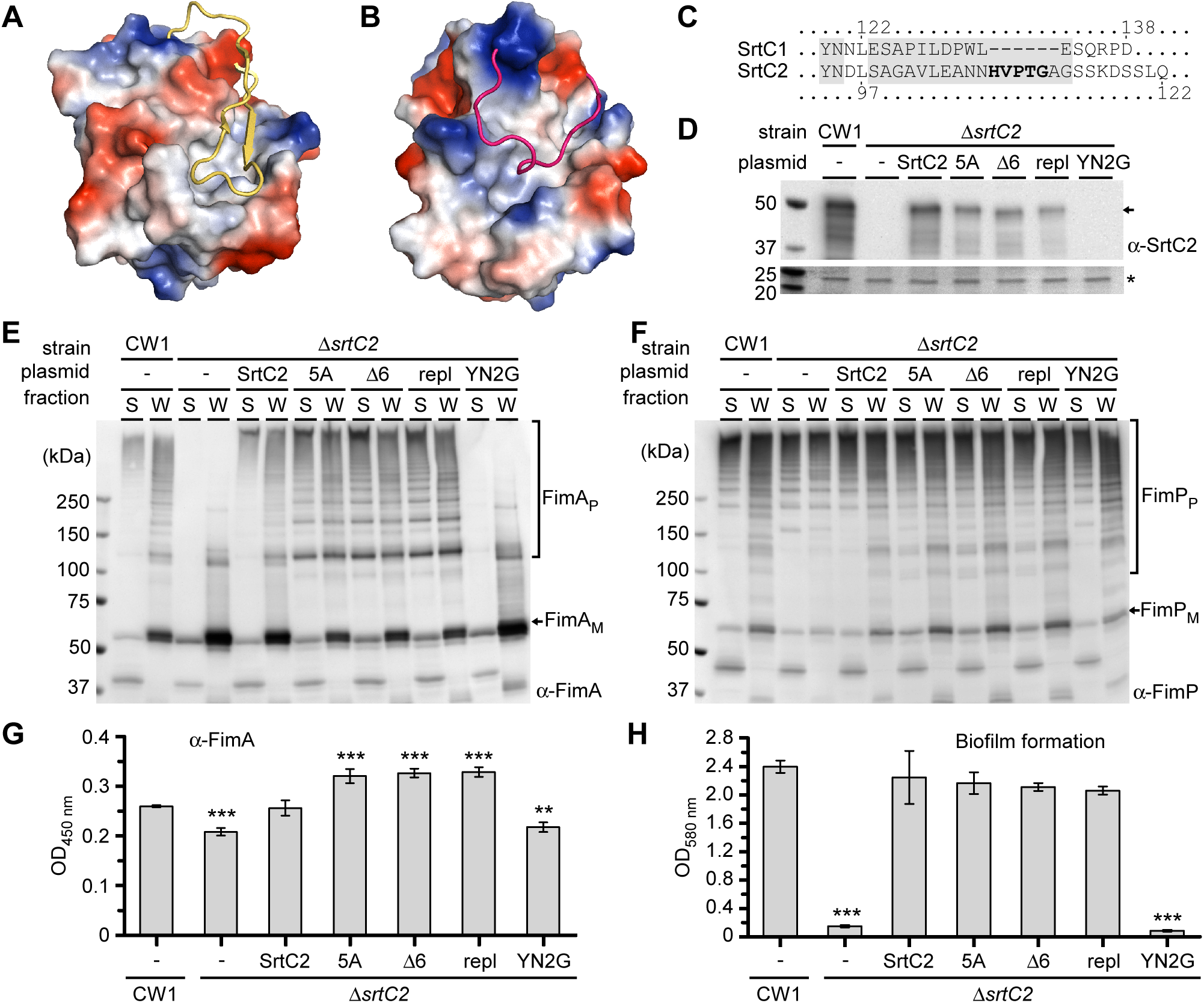
The lid is dispensable for SrtC2-catalyzed polymerization of FimA pilins. **(A-B)** The lid of SrtC2 (yellow) and SrtC1 (red) is displayed as cartoons over the electrostatic surface potential of the sortase score structures, with red for negative charge and blue positive). **(C)** Shown is a sequence alignment between the SrtC1 and SrtC2 lids, with numbers marking the lid region of each enzyme and 5 bolded letters indicating residues for alanine-substitution. **(D)** The following strains were used for cell fractionation; parent strain (CW1), its isogenic mutant Δ*srtC2* and this mutant expressing wild-type SrtC2 (pC2) or its variants, with alanine-substitution of 5 bolded residues in (C) (5A), deletion of residues HVPTGA (Δ6), the lid region (residues 98-114) replaced with the SrC1 lid (residues 123-133) (repl), or residues YN mutated to G (YN2G). Protein samples from the membrane fractions were immunoblotted with antibodies against SrtC2 (α-SrtC2), with a non-specific band (asterisk) used as loading control. **(E-F)** Protein samples from the culture supernatant (S) and cell wall (W) fractions were analyzed by immunoblotting with antibodies against FimA (α-FimA) (E) or FimP (α-FimP) (F). Pilus polymers (P), monomers (M), and molecular weight markers (kDa) are specified. **(G)** Surface display of FimA of indicated strains was determined by whole-cell ELISA. **(H)** Biofilms of indicated strains were cultivated and quantified by crystal violet measurement. Results in G-H are presented as average of three independent experiments performed in triplicate; ***, P< 0.001; **, P< 0.01.

Based on DALI analysis (28), the closest structure to the SrtC2 is *A. oris* SrtC1 (PDB:2XWG) (19), having Z-score and RMSD values of 23.8 and 2.1 for 162 structurally equivalent residues, respectively (Fig. 1C), together with both proteins sharing 42% of amino acid identity. Nonetheless, the lid of SrtC2 is longer than that of SrtC1, which consists of a single 17-residue strand blocking the catalytic triad (Fig. 1B and Fig. 3A-3C). In addition to the aforementioned unique lid conformation and β11-strand, another major difference in the SrtC2 structure compared to that of SrtC1 lies in the two loops made of residues 223-232 and 246-255. Specifically, the latter’s conformation is critical for the enzyme activity due to the presence of two key active site residues, C246 and R255. The shift of the loop causes a change, in a relative position of the SrtC2 catalytic triad (Fig. 1D), resulting in the C246 C-α atom moving about 4 Å towards the H184 residue. All closest SrtC2 structural homologs have preserved positions of catalytic triad residues. Also, the loops comprising catalytic residues have very similar conformations (Fig. 1E). Potentially, the unique loop conformation in the SrtC2 structure might be forced by contacts from neighbor molecules inside crystals. Nevertheless, it shows potential for flexibility of these loops in the protein itself.

We also compared the SrtC2 structure to that of SrtA, a class E sortase that had been reported previously (PDB:5UTT) (18). Remarkably, despite of low amino-acid sequence identity of 17.5%, both structures share significant structural similarity with RMSD equal to 2.4 Å. Moreover, the conserved sortase core is very similar in both structures. Yet, it is important to recall that SrtA as a housekeeping sortase has no flexible lid to cover its active site. The N-terminus of SrtA, corresponding to the N-terminal part of SrtC2 which includes a lid region, is unstructured and positioned on the protein surface area matching the SrtC2 α1 helix location (Fig. 1F). Curiously, the N-terminus contains several conserved residues, including the YN motif (Fig. 2). Additionally, both structures significantly differ in the same regions mentioned above in SrtC2-SrtC1 structure comparison.

### Mutations of the SrtC2 lid do not affect assembly of type 2 fimbriae and biofilm formation

Given the major differences found in the lid of SrtC2 as compared to that of SrtC1 reported above, as well as the features of the SrtC2 N-terminus relative to the unstructured domain of SrtA (Fig. 1-2 and Fig. 3A-3B), it is logical to investigate whether targeted mutations altering these features would affect SrtC2 activity and substrate specificity or not. We, therefore, generated recombinant plasmids expressing SrtC2 with the following mutations: (i) the conserved residues YN mutated to G (YN2G), (ii) the HVPTG sequence of the lid changed to five Ala (5A), (iii) deletion of this extended sequence (Δ6), or (iv) a SrtC2 variant with its lid segment from S98 to G114 replaced by the equivalent lid segment of SrtC1 (residues E123 to E133) (repl). The wild-type and mutant plasmids were introduced into an *A. oris* mutant devoid of *srtC2* (Δ*srtC2*). To first examine if these mutations alter protein stability, we subjected these strains to cell fractionation and performed western blotting of membrane cell lysates using an antibody against SrtC2 as previously reported (29). As compared to the parent strain (CW1) or the Δ*srtC2* mutant expressing wild-type SrtC2 (pC2 or pSrtC2), the Δ*srtC2* mutants expressing SrtC2 with 5A, Δ6, or repl mutation showed some reduction of SrtC2 protein levels, whereas no SrtC2 was observed in the YN2G mutant (Fig. 3D), indicating that the mutation of the conserved YN motif affects SrtC2 stability.

Upon establishing the relative protein levels of the mutant enzymes, we next investigated whether specific mutations affect function, specifically the formation of type 2 fimbriae required for biofilm formation (11,30). In this experiment involving subcellular fractionations, the supernatant (S) and cell wall (W) fractions prepared from the above strains were subjected to SDS-PAGE and subsequent immunoblotting with antibodies against the major shaft proteins FimA (α-FimA) and FimP (α-FimP). Notably, as determined from the relative sizes and intensities of the pilus polymers displayed on PAGE, the lid mutants showed no significant defect in pilus polymerization as compared to the parent strain (CW1) and the wild-type *srtC2*-complementing strain (Δ*srtC2*/pSrtC2), despite some reduction in the protein levels of the mutant enzymes; however, as expected from the lack of SrtC2, the YN2G mutant did not form FimA polymers, similar to the Δ*srtC2* mutant (Fig. 3D and 3E). By contrast, it is important to emphasize that the assembly of type 1 fimbriae (FimP), which requires SrtC1 activity, was not altered at all by any one of the above mutations (Fig. 3F). To quantify how much of the polymerized pili made to the bacterial cell wall and displayed at the surface, we performed whole-cell ELISA with α-FimA, which revealed that the lid mutations significantly increased the surface expression of FimA; by comparison, the YN2G mutant was defective in type 2 fimbrial assembly, as expected, similar as the Δ*srtC2* mutant (Fig. 3G). Furthermore, the lid mutants produced biofilms at levels comparable to that of the parent strain, while the YN2G mutant was unable to form biofilms (Fig. 3H).

Next, it was important to visualize the quantity and quality of the cell surface fimbriae directly. To do so, we analyzed the same set of bacterial strains described above by immuno-electron microscopy (IEM) (18), whereby *A. oris* cells were immobilized on carbon-coated nickel grids, stained with polyclonal antibodies against the two fimbrillins FimA or CafA. Consistent with the above western blot and ELISA data, no significant alterations in surface assembly of fimbriae were observed in the lid mutants as compared to the parent strain (Fig. 4A-4K and Fig. 4M-4O), while the phenotype of the YN2G mutant SrtC2 mirrored the Δ*srtC2* mutant (Fig. 4L and 4P).

**Figure 4:**
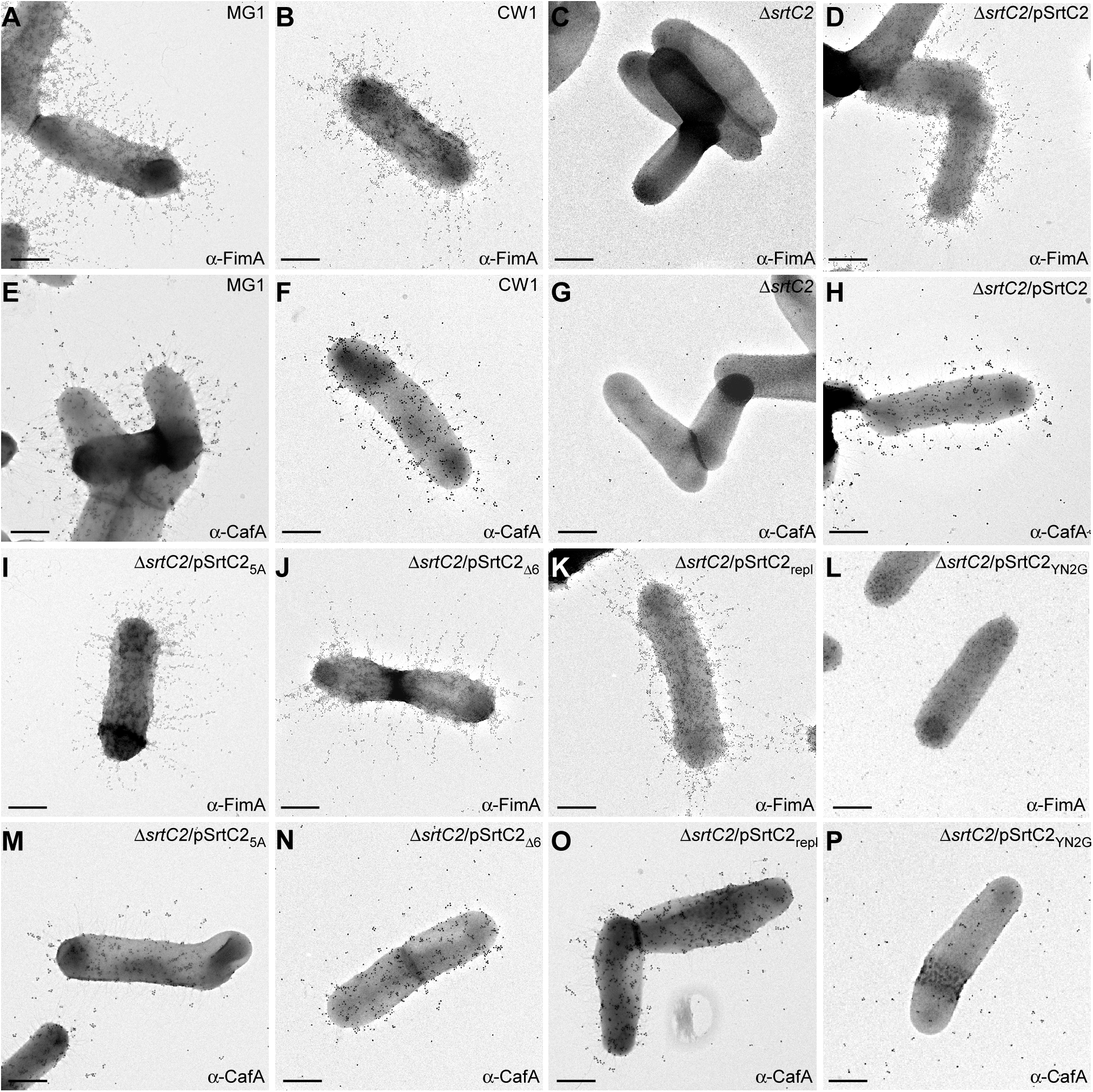
Immunoelectron microscopic analysis of pilus assembly. *A. oris* cells of indicated strains were analyzed by immunoelectron microscopy with specific anti-sera, α-FimA **(A-D; I-L)** and α-CafA **(E-H; M-P)**, followed by goat anti-rabbit IgG conjugated to 12-nm and 18-nm gold particles, respectively; scale bars of 0.5 μm.

Curiously, the lid mutant strains appear to produce less pilus tip CafA than the parent and the SrtC2-complementing strain (Fig. 4, compare panels E and F with panels M-O), and CafA was released in the surroundings with the YN2G mutation similar to what is observed with the Δ*srtC2* mutant (Fig. 4, compare panel G with panel P). Altogether, the results indicate that mutations of the lid do not alter the polymerizing activity of SrtC2 involving the major fimbrial shaft FimA and normally displays the surface fimbriae that are required for normal biofilm formation.

### SrtC2 lid mutations diminish surface display of CafA and CafA-mediated polymicrobial interaction

Since SrtC2-catalyzed assembly of CafA in the type 2 fimbriae mediates interspecies interaction or coaggregation (11,14,30), we next characterized the above mutant strains in previously established coaggregation assays using streptococci, given the apparent reduction of CafA with the lid mutants as mentioned above (Fig 4). Briefly, *A. oris* cells were mixed with *Streptococcus oralis* (So34) in equal volumes and coaggregation was determined both optically (Fig. 5A) (14) and quantitatively (Fig. 5B) (31). Compared to the *A. oris* cells expressing WT SrtC2 (CW1 and Δ*srtC2*/pSrtC2), each strain with the lid mutations showed severe defects in coaggregation, while the YN2G mutant SrtC2 was unable to aggregate with So34 similar to the Δ*srtC2* mutant (Fig. 5A and Fig. 5B). To directly examine whether the lid mutations affect the incorporation of CafA as the pilus tip, we analyzed the supernatant (S) and cell wall (W) fractions of the same set of strains by SDS-PAGE and immunoblotting with antibodies against CafA (α-CafA). As shown in Fig. 5C, no apparent defects of CafA pilus incorporation were observed in these mutant strains, except for the YN2G mutant as expected (Fig. 5C) considering its defective expression of SrtC2 (Fig. 3D). However, we noticed that the lid mutant strains appeared to display reduced amounts of cell wall-anchored pilus tip CafA than the SrtC2-complementing strain (Fig. 5, compare the W lane of pSrtC2 with that of 5A, Δ6, and repl). This is entirely consistent with the phenotypes of the mutants we detected by immuno-electron microscopy as mentioned above.

**Figure 5:**
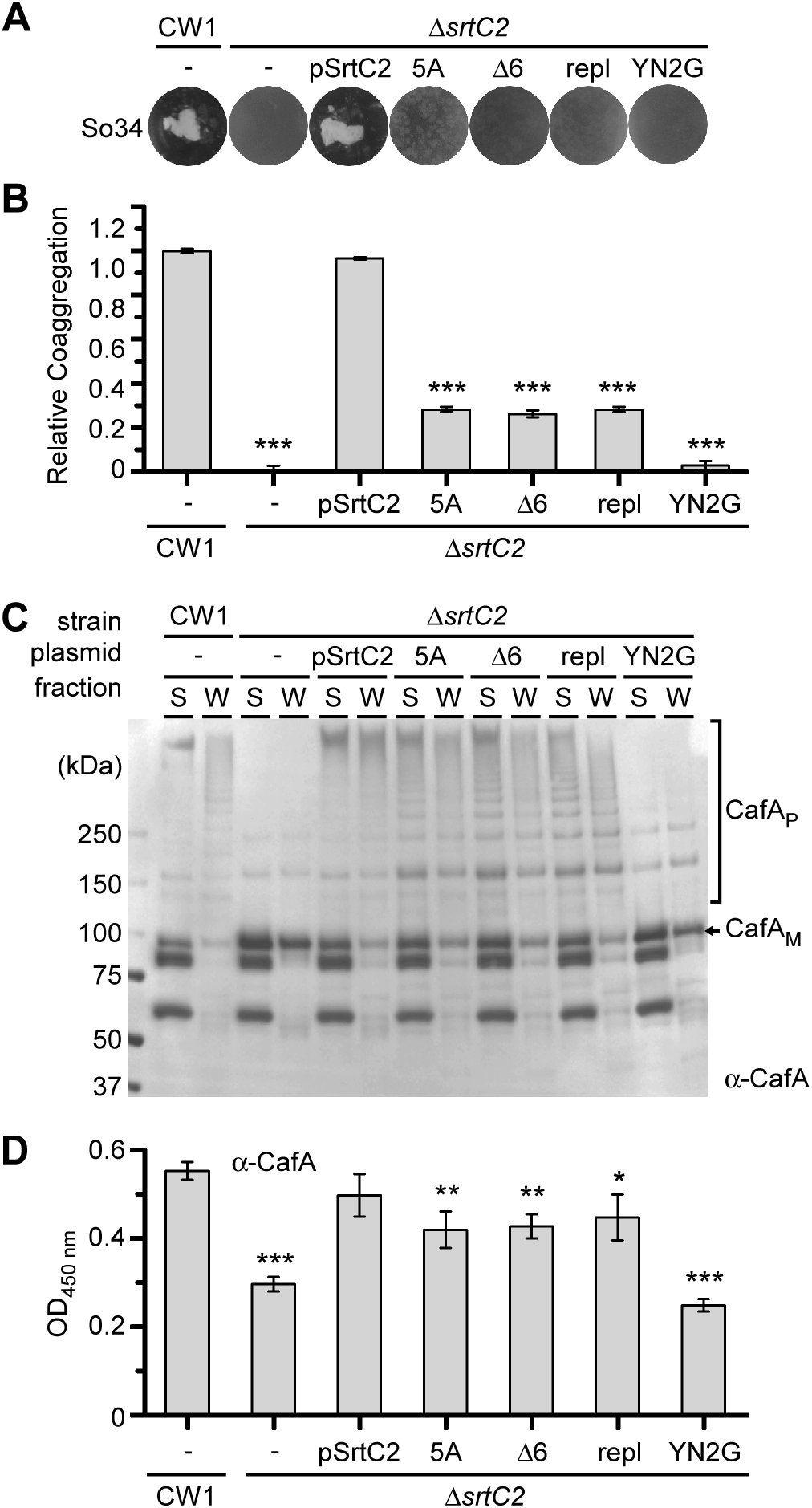
Mutations of the SrtC2 lid affect bacterial coaggregation and CafA surface display. **(A-B)** Interaction between *S. oralis* So34 and indicated *A. oris* strains was determined by a visual coaggregation assay (29) (A) or quantitatively (B). **(C)** Similar to the experiments in Fig. 3E-3F, protein samples from the culture supernatant (S) and cell wall (W) fractions of indicated strains were immunoblotted with antibodies against CafA (α-CafA). CafA monomers (M), polymers (P) and MW makers are shown. (D) Whole cell ELISA was performed with indicated strains to quantify the cell surface display of CafA. (***, P< 0.001; **, P< 0.01; *, P< 0.05).

To further scrutinize this observation, we quantified the protein level of surface displayed CafA by whole-cell ELISA with α-CafA, and indeed we found that all mutants expressed significantly less CafA on the bacterial surface, with most severe defect noted for the YN2G mutant (Fig. 5D). Therefore, we conclude that the coaggregation defect observed in the lid mutants is due to the reduced level of surface displayed CafA caused by mutations of the SrtC2 lid.

### Specific mutations of the SrtC2 lid alter cell wall anchoring activity for the major shaft pilins

To critically examine the cell anchoring function of SrtC2 in the absence of SrtA, we used a previously generated triple mutant strain which is devoid of SrtC2 and SrtA, as well as the major surface glycoprotein GspA, mutations of which suppress the lethality of *srtA* deletion in *A. oris* (23). As we reported previously, this triple mutant strain is coaggregation-negative, unable to produce FimA polymers, and is severely defective in cell wall anchoring of FimP polymers (see also Fig 6A-6C; first two circles/lanes). Ectopic expression of wild-type SrtC2 in this mutant strain restored FimA pilus polymerization, and to some degree, the cell wall anchoring of FimA polymers (Fig. 6B; lanes SrtC2). Note that the extended FimA pili synthesized and assembled on cell surface failed to mediate coaggregation (Fig. 6A; circle SrtC2), an observation that agrees with our previous report (14). By comparison to wild-type SrtC2, the alanine-substitution of the lid did not affect FimA pilus polymerization or cell wall anchoring (Fig. 6B; lanes SrtC2_5A_). Moreover, consistent with the results presented in Fig. 3E, partial deletion of the SrtC2 lid did not alter FimA pilus polymerization, yet cell wall anchoring of FimA polymers was significantly reduced in this case (Fig. 6B; lanes SrtC2_Δ6_). Similarly, swapping the lid of SrtC2 with that of SrtC1 also reduced cell wall anchoring of FimA polymers, though the polymerization of FimA pilins was not affected (Fig. 6B; last two lanes).

**Figure 6:**
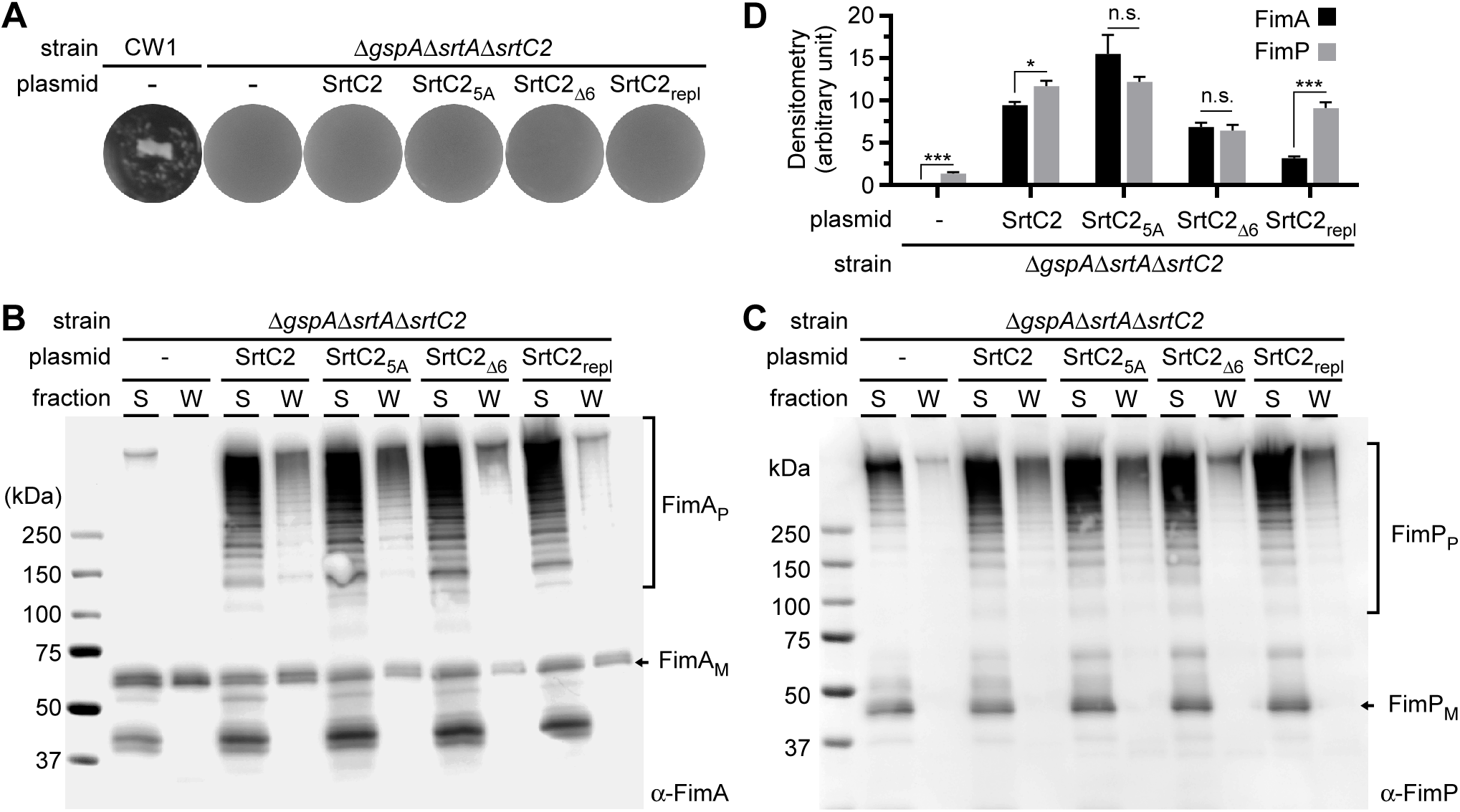
Mutations of the SrtC2 lid alter cell wall-anchoring specificity for the major pilins. **(A)** Coaggregation between *S. oralis* So34 and indicated *A. oris* strains was determined by a visual coaggregation assay. **(B-C)** Similar to the experiments in Fig. 3E-3F, protein samples from the culture supernatant (S) and cell wall (W) fractions of indicated strains were immunoblotted with α-FimA (B) or α-FimP (C). Pilin monomers (M), polymers (P) and MW makers are shown. **(D)** The intensity of cell wall fractions in (A) and (C) was quantified by densitometry using ImageJ. The results are presented as average of three measurements with standard deviations; ***, P< 0.001; **, P< 0.01; *, P< 0.05, and n.s., P>0.05.

To examine the effects of the aforementioned mutations of the SrtC2 lid on cell wall anchoring of FimP polymers catalyzed by SrtC1, we immunoblotted the same set of samples described above with α-FimP (Fig 6C). In line with our previous study (18), ectopic expression of wild-type SrtC2 in the triple mutant strain enhanced the cell wall anchoring of FimP polymers (Fig. 6C; compare first two lanes with lanes SrtC2). Alanine-substitution of the SrtC2 lid did not change cell wall anchoring of FimP polymers (Fig. 6C; lanes SrtC2_5A_); however, partial deletion of the SrtC2 lid significantly decreased this process (Fig. 6C; lanes SrtC2_Δ6_). Most remarkably, swapping the lid of SrtC2 with that of SrtC1 enhanced the cell wall anchoring of FimP polymers as compared to the SrtC2_Δ6_ mutant (Fig. 6C; last two lanes). Consistently, quantification of cell wall anchoring of FimA and FimP polymers in the aforementioned strains by densitometry showed similar results as the above (Fig. 6D).

To further confirm the distinctive pilus assembly phenotypes reported above, we employed IEM to analyze the same set of strains (Fig. 7). Compared to the parent strain, the triple mutant was not able to produce FimA pili (Fig. 7; compare panel A with B) and displayed an inefficiently assembled FimP polymers on the cell surface (Fig. 7; compare panel H with I). As expected, ectopic expression of wild-type SrtC2 promoted assembly of extended FimA and FimP pili (Fig. 7; panels C with J). Consistent with our immunoblotting results, no apparent defects in surface assembly of FimA and FimP pili were observed in the SrtC2_5A_ mutant as compared to the pSrtC2 strain (Fig. 7; panels D and K). Significantly, both partial deletion of the SrtC2 lid and swapping of this lid with that of SrtC1 reduced surface assembly of FimA pili (Fig. 7; panels E and F). While partial deletion of the SrtC2 lid caused some secretion of FimA pili, and their reduced surface assembly, lid-swapping mutation enhanced surface assembly of FimP pili (Fig. 7; compare panel L with panel M).

**Figure 7:**
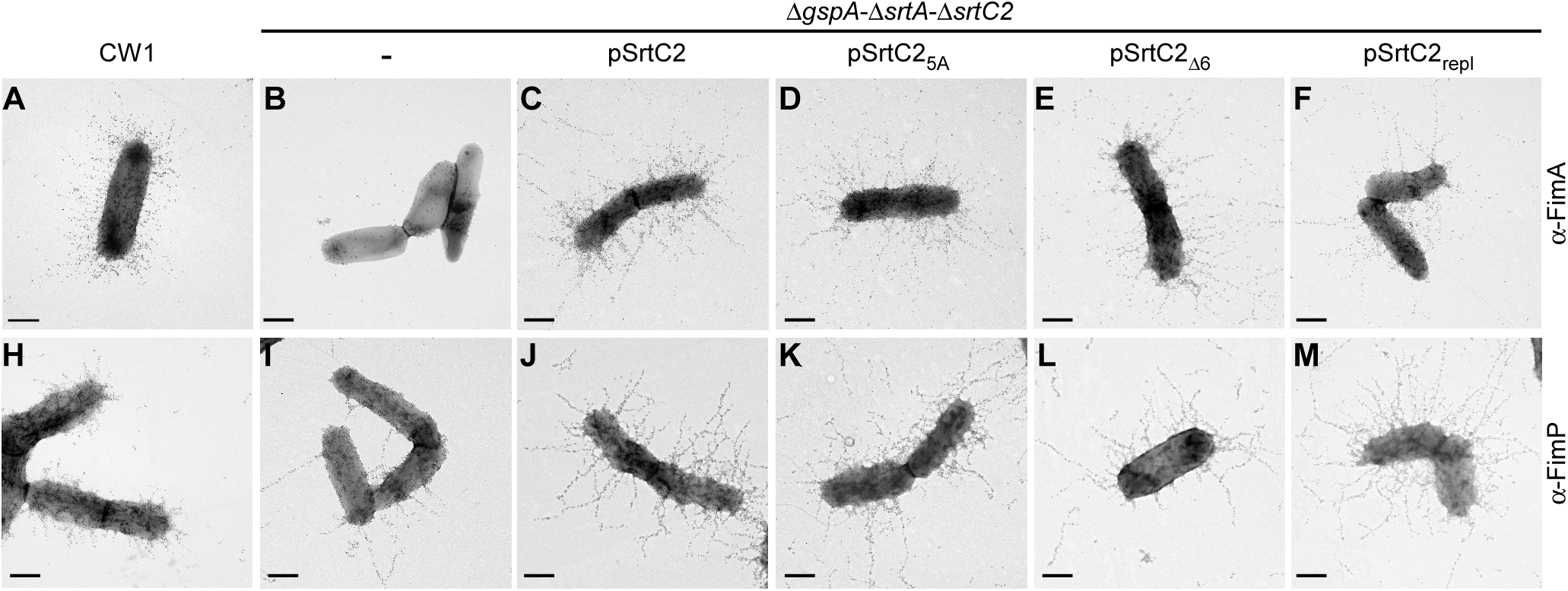
Immunoelectron microscopic analysis of pilus assembly. *A. oris* cells of indicated strains were analyzed by immunoelectron microscopy with specific anti-sera, α-FimA **(A-F)** and α-FimP **(H-M)**, followed by goat anti-rabbit IgG conjugated to 12-nm; scale bars of 0.5 μm.

To quantify the level of pilus surface display, we adapted a previously published method of immunofluorescence to probe cell wall anchoring of Gram-positive bacterial pili (32), whereby the fluorescent signal of mid-log phase cells labeled with AlexaFluor Plus 594 via α-FimA (Fig. 8A) or α-FimP (Fig. 8C) was quantified by ImageJ. Consistent with the above results (Fig. 6-7), the SrtC2 mutant enzyme with partial deletion of its lid (SrtC2_Δ6_) reduced cell wall anchoring of FimA (Fig. 8B) and FimP polymers (Fig. 8D), as compared to the wild-type enzyme, whereas the SrtC2 enzyme with its lid replaced by the lid of SrtC1 (SrtC2_repl_) was deficient in cell wall anchoring of FimA polymers while increasing cell wall anchoring of FimP polymers, relative to the SrtC2_Δ6_ enzyme (Fig. 8B & 8D). Collectively, the simplest interpretation of these results is that the lid region of SrtC2 is an important structural element that governs and modulates the cell wall anchoring activity of this enzyme.

**Figure 8:**
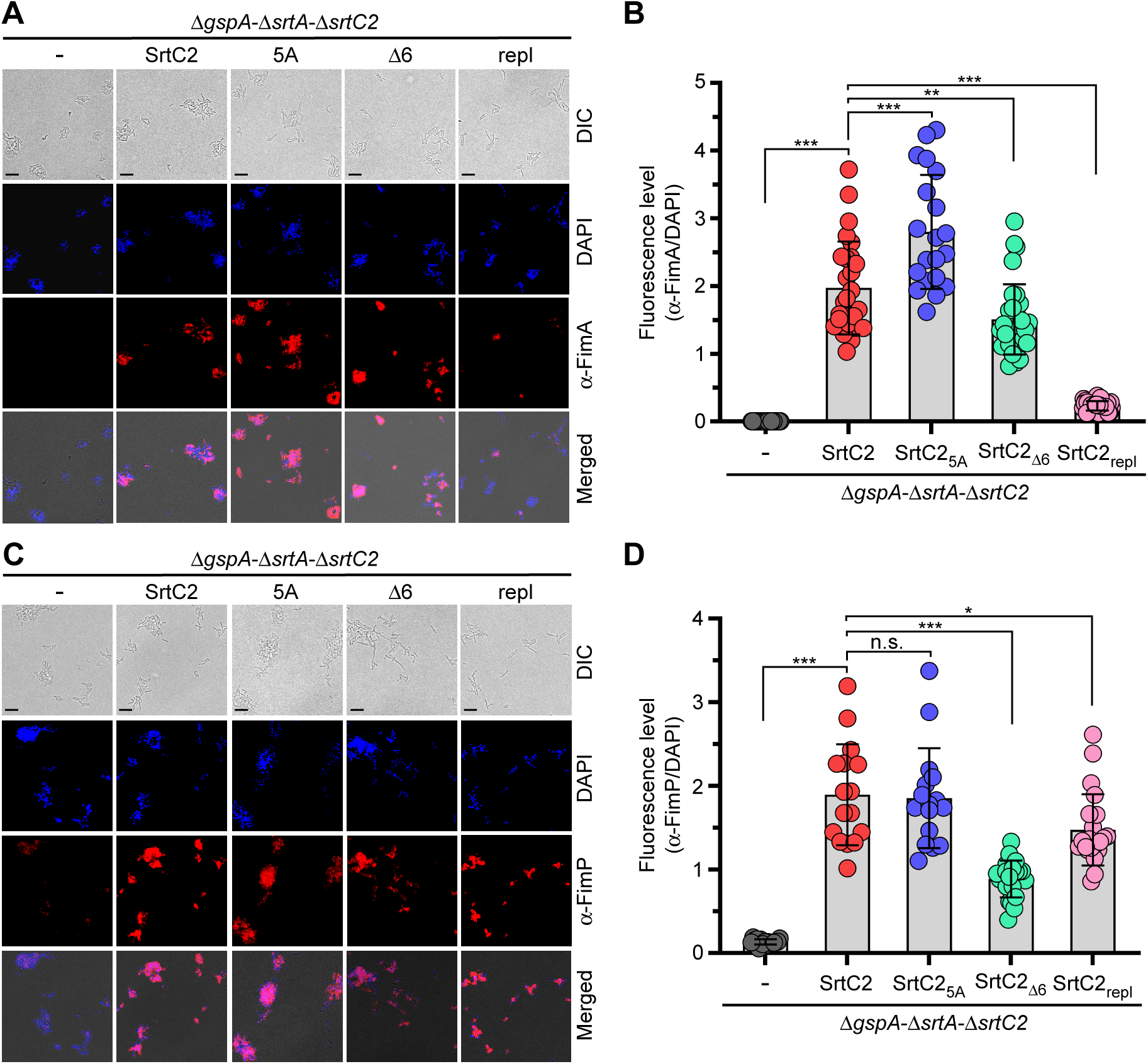
Immunofluorescence microscopy of pilus surface display. **(A-D)** Mid-log phase cells of indicated strains were fixed and stained specific antibodies α-FimA (A) or α-FimP (C), followed by AlexaFluor Plus 594 (red), with DAPI used for normalization. For quantification, the results are presented as averaged fluorescence levels of FimA (B) or FimP (D) signal, relative to DAPI signal. Significant analysis was performed by one-way ANOVA with GraphPad Prism; *, P<0.05; **, P<0.01; ***, P<0.001, and n.s., P>0.05.

## DISCUSSION

Numerous studies have established that the cell wall anchoring of covalently linked pilus polymers in Gram-positive bacteria requires the enzymatic activity of the conserved housekeeping sortase (either class A or class E family) (15,21). It follows that the genetic disruption of the housekeeping sortase would fail to join pili to the cell wall and consequently cause pilus polymers to be excreted into the extracellular milieu, as we demonstrated in the case of *C. diphtheriae* when its housekeeping sortase *srtF* gene is deleted (32). Contrary to this logical scenario was our observation in *A. oris* that the deletion of its housekeeping sortase *srtA* did not obliterate the cell surface assembly of certain pili, but rather caused the assembly of extensively long pilus polymers and their anchoring to the bacterial cell wall (23). The mysterious activity for cell wall display of pilus polymers without the classical anchoring sortase in this case turned out to be the very sortase SrtC2 that polymerizes type 2 fimbrillins. By biochemical and genetic analyses, we discovered that the pilus-specific sortase SrtC2 not only polymerizes the cognate FimA pilins, linking the FimA polymers to the CafA or FimB tip pilins, but it also catalyzes the subsequent cell wall anchoring of the FimA polymers and the non-cognate FimP polymers in the absence of SrtA – the sortase dedicated to link many proteins the cell wall rather promiscuously (18). By a combination of structural, genetic, biochemical and cytologic studies reported here, we unveiled the structural basis for this unique enzymatic innovation: how a sortase has evolved to carry out both pilus polymerization and cell wall anchoring functions of sortases by utilizing a simple ‘lid’ structure to modulate the catalytic core of the enzyme yet still maintaining exquisite specificity for its substrates.

By X-ray crystallization, we describe here a well-resolved 3-D structure of this dual function sortase (SrtC2) that harbors a canonical sortase fold and a structural lid typical of class C sortase enzymes (Fig. 1). Intriguingly, the lid sequence of SrtC2 is longer than that of another pilus-specific sortase SrtC1 of the organism (Fig. 2 and Fig. 3C). This led us to consider that the extra sequence might provide SrtC2 with the differential activity and specificity displayed by this enzyme. To test this hypothesis, we generated several SrtC2 variants – alanine-substitution of this extra sequence (SrtC2_5A_), deletion of this extra sequence (SrtC2_Δ6_) and swapping the lid of SrtC2 with that of SrtC1 (SrtC2_repl_), as well as glycine-substitution of the YN residues at the N-terminus of SrtC2 (SrtC2_YN2G_). Although the mutations caused some reduction in the protein levels of the mutant enzymes except for YN2G mutation that caused protein instability, each of the other mutations did not abrogate but rather enhance the polymerization activity of SrtC2, (Fig. 3 and Fig. 4). This is consistent with the conclusion from previous studies of other pilus-specific sortases that the lid itself is dispensable for pilus polymerization (33,34). The increased FimA polymerization and surface display by mutations of the lid might be due to increased accessibility of FimA substrates for the catalytic center as similarly observed with the lid mutations in the pilus-specific sortase SrtA of *C. diphtheriae* (34).

An initial hint for the role of the lid in substrate specificity came from an immunoelectron microscopy (IEM) experiment that monitored surface assembly of CafA and a robust functional assay that measured pilus-mediated polymicrobial interaction (or coaggregation); all of the stably expressed lid mutants exhibited a reduced level of surface CafA (Fig. 4, compare panel F with panels M-P; Fig. 5, panels C-D), paralleled by significant defects in bacterial coaggregation with one of the natural partners, *S. oralis* (Fig. 5A-5B). Because CafA incorporation into the FimA polymers is the first step of pilus polymerization reactions (16) and CafA has a slightly different CWSS (the LPXTG motif, i.e. IPFTG (14) than that of FimA and FimP having LPLTG), it seems plausible that the lid mutations negatively affected the specificity of SrtC2 toward CafA. Notably, this is the first time that a structural determinant specifically involved in substrate recognition of sortase SrtC2 has been unveiled.

To examine the cell wall anchoring function of SrtC2, we utilized a mutant strain that lacks both *srtA* and *srtC2*, as well as *gspA* coding for a major cell wall anchored glycoprotein whose absence suppresses the lethality of *srtA* deletion (23). In this triple mutant strain, expression of the SrtC2_5A_ mutant allows normal pilus polymerization and cell wall anchoring of both FimA and FimP polymers, compared to complementation by wild-type SrtC2 (Fig. 5-8). By contrast, deletion of the extra sequence of SrtC2 and lid swapping with the equivalent sequence from SrtC1 (another pilus-specific sortase) reduce cell wall anchoring of FimA polymers, whereas these same mutations enhance the cell wall anchoring of FimP polymers (Fig. 5-8). Given the differential activity of cell wall anchoring of FimA and FimP polymers with these lid mutations, while the polymerization activity of the mutant enzymes is maintained, the reduced protein levels of these mutant enzymes cannot fully account for their differential cell wall anchoring activities for both FimA and FimP polymers. Therefore, the results suggest that the conformation created by the extra sequence of the SrtC2 lid modulates substrate specificity for SrtC2-mediated transpeptidation reactions that give rise to pilus polymerization or cell wall anchoring.

Normally, pilus polymerization is terminated by the housekeeping SrtA, which links the terminal polymer to the cell wall. A key question arises as to how pilus polymerization is terminated when SrtA is absent and cell wall anchoring is mediated by SrtC2. Since *A. oris* fimbriae are heterodimeric, the last subunit of pilus polymers, i.e. the terminal shaft pilin subunit, serves as the pilus base that anchors to the cell wall. We suspect that the net availability of substrate pilins, i.e. FimA monomers, at the catalytic center will dictate whether the pilus continues to extend or gets terminated and anchored to the cell wall. According to this scenario, SrtC2 continues to polymerize FimA pilins to elongate the pilus structure until the FimA pilins are exhausted at the assembly center. When the FimA pilin is not available, the nucleophiles provided by the lipid II precursors resolve the acyl enzyme intermediate with the last FimA molecule, thereby terminating pilus polymerization and permitting cell wall anchoring of pilus polymers.

Another intriguing question is how SrtC2 mediates cell wall anchoring of FimP polymers and whether SrtC2 anchors other SrtA substrates to the cell wall. It is likely that the last FimP molecule forming the acyl enzyme intermediate with SrtC2 is attached to peptidoglycan in a similar manner as the cell wall anchoring process for FimA polymers. Given that FimA and FimP have the same LPLTG motif, another possibility is that a FimA-SrtC2 intermediate may terminate FimP polymerization, linking FimP polymers to peptidoglycan with FimA as the last subunit. While future experiments will explore these possibilities, the finding of the dual activity of SrtC2 provides an intriguing new insight into the evolution of sortase enzymes that may have important implications in developing new tools for sortase-mediated bioengineering.

## EXPERIMENTAL PROCEDURES

### Bacterial strains, plasmids, and cell culture

Bacterial strains and plasmids used in this study are listed in Table 2. *Actinomyces* strains were grown in heart infusion broth (HIB) or on HIB agar plates at 37°C. Streptococci were grown in Brain Heart Infusion (BHI) supplemented with 1% glucose in an anaerobic chamber. When required, kanamycin was added into medium at a final concentration of 50 µg ml^-1^.

### Plasmid construction

To generate recombinant plasmids expressing SrtC2 and its variants, a DNA segment encompassing the *srtC2* promoter region and *srtC2* gene in pJRD215 (8) was subcloned into pCWU10 (31) at *SalI* and *HindIII* sites to generate pSrtC2. This recombinant plasmid was used as template to generate SrtC2 variants by site-directed mutagenesis as previously reported (18). Briefly, a pair of phosphorylated primers harboring designed mutation(s) (Table 3) were used to amplify a linearized DNA segment, which was purified and circularized by T4 ligase. The circularized plasmids generated were transformed into *E. coli* DH5α, and the sequence of each *srtC2* mutant was confirmed by DNA sequencing before being introduced into *A. oris* by electroporation.

For generation of a recombinant vector expressing His-tagged SrtC2, a DNA fragment coding for a SrtC2 molecule spanning residues 60 to 283 was cloned into pMCSG7 using ligation-independent cloning as previously reported (35). The generated plasmid (pMSCSG7-SrtC2) (Table 2) was transformed in *E. coli* BL21 (DE3) for protein expression.

### Protein purification for crystallization

*E. coli* cells harboring pMSCSG7-SrtC2 were cultured in LB medium supplemented with ampicillin (100 µg/mL) at 37°C. When the optical density at 600 nm reached 0.8, cultures were transferred to 4°C for 1 hour. Isopropyl β-D-1-thiogalactopyranoside (IPTG) was added to a final concentration of 0.5 mM for overnight induction at 18°C. Cells were harvested by centrifugation, disrupted by sonication, and the insoluble cellular material was removed by centrifugation. SrtC2 protein was purified using Ni-NTA (Qiagen) affinity chromatography with the addition of 5 mM β-mercaptoethanol in all buffers. The protein was digested with 0.15 mg TEV protease per 20 mg of purified SrtC2 protein for 16 h at 4°C, and then passed through a Ni-NTA column to remove both the TEV protease and cleaved N-terminal tags. The final step of purification was performed with size-exclusion chromatography on HiLoad 16/60 Superdex 200pg column (GE Healthcare) in 25 mM HEPES buffer pH 7.5, 200 mM NaCl and 1 mM DTT. The protein was concentrated on Amicon Ultracel 10K centrifugal filters (Millipore) to 23 mg/ml concentration.

### Protein crystallization

The initial crystallization condition was determined with a sparse crystallization matrix at 4°C and 16°C temperatures using the sitting-drop vapor-diffusion technique using MCSG crystallization suite (4 screens) (Microlytic), Pi-minimal and Pi-PEG screens (Jena Bioscience) (36). The best crystals were obtained after 2 months of incubation at 4°C temperature from the A3 conditions (20% PEG 8000, 0.2 M sodium chloride, 0.1 M disodium phosphate: citric acid, pH 4.2). Crystals selected for data collection were briefly soaked in crystallization buffer with addition of 15% ethylene glycol as cryo-protectant and then flash-cooled in liquid nitrogen.

### Data Collection, Structure Determination and Refinement

Single-wavelength x-ray diffraction data were collected at 100 K temperature at the 19-ID beamline of the Structural Biology Center at the Advanced Photon Source at Argonne National Laboratory using the program SBCcollect (37). The intensities were integrated and scaled with the HKL3000 suite (38). The SrtC2 protein structure was determined by molecular replacement using HKL3000 suite incorporating MOLREP program (39) and *A. oris* sortase SrtC1 (PDB: 2XWG) (19) as the starting model. Several rounds of manual adjustments of structure models using COOT (40) and refinements with Refmac program (41) from CCP4 suite were performed. The stereochemistry of the structure was validated with PHENIX suite (42) incorporating MOLPROBITY (43) tools. A summary of data collection and refinement statistics is given in Table 1.

### Whole cell enzyme-linked immunosorbent assay (ELISA)

The level of FimA and CafA displayed on the bacterial cell surface in different strains was determined by whole cell ELISA according to a published procedure (18). In brief, log-phase cells were harvested and washed with PBS before being resuspended in 0.1 M bicarbonate buffer, pH 9.6 and the cell densities were normalized to OD_600_ of 1.0. 100 μl-aliquots were used to coat high-binding 96-well plates (Corning Costar) for 1 h at 37°C prior to washing and blocking with 2% bovine serum albumin (BSA) in phosphate-buffered saline (PBS). To detect surface proteins, coated cells were treated with appropriate antibodies (α-FimA, 1:5000, α-CafA, 1:1000) for 2 h, followed by horseradish peroxidase (HRP)-conjugated anti-rabbit IgG (1:10000) and 3,3′,5,5′-tetramethylbenzidine (Invitrogen). The reactions were quenched by addition of 1N H_2_SO_4_. Absorbance at 450 nm was measured using a Tecan M1000 plate reader.

### Cell fractionation and Western blotting

Cell fractionation and Western blotting were followed according to a published protocol (29). Briefly, mid-log phase cultures of *A. oris* strains normalized to equal cell density (OD_600_ of 1) were subjected to cell fractionation, and protein samples from the culture medium (S) and cell wall (W) fractions were obtained by precipitation with 7.5% trichloroacetic acid (TCA), followed by washing with cold acetone. Protein samples were dissolved in sodium dodecyl sulfate (SDS) sample buffer containing 3M urea, separated by 3-12% Tris-glycine gradient gels, and subjected to immunoblotting with specific antisera (α-FimA at 1:20000 dilution and α-CafA at 1:4000).

### Bacterial coaggregation assay

Coaggregation between *A. oris* and *S. oralis* So34 was performed as previously described (31). Briefly, stationary-phase cultures of bacterial strains grown in CAMG complex medium with 0.5% glucose were harvested by centrifugation. Bacterial cells were washed in Tris-buffered saline (TBS, pH 7.5) containing 0.1 mM CaCl_2_ and normalized to an OD_600_ of 2.0 (approximately 2 x 10^9^ CFU/ml). 0.5-ml aliquots of *Actinomyces* and streptococcal cell suspensions were mixed in 24-well plates for a few minutes on a rotator shaker and the coaggregation were imaged by Alpha Imager (ProteinSimple). For quantification, 1-mL aliquots of coaggregation mixtures were subjected to centrifugation at 100 x g for 2 min. The optical density (OD_600nm_) of the obtained supernatants was measured to determine the coaggregation efficiency.

### Immunoelectron microscopy (IEM)

Overnight grown cells of various *A. oris* strains on HIA plates were washed once and suspended in PBS. A drop of bacterial suspension was placed onto carbon-coated nickel grids. Bacterial cells on grids were first stained with specific antibodies against *A. oris* pilins (α-FimA, 1:100; α-FimP, 1:100; and α-CafA, 1:50), followed by incubating with the secondary antibody conjugated with gold particles (12 nm for FimA and 18 nm for CafA) and staining with 1% uranyl acetate before imaged under an electron microscope.

### Immunofluorescence microscopy

To determine the levels of cell wall anchoring of pilus polymers, immunofluorescence was employed based on previously published protocols with some modification (32,44). Briefly, mid-log phase cultures of *A. oris* strains normalized to equal cell density (OD_600_ of 1), and harvested cells were washed with and resuspended in PBS to OD_600_ of ∼ 0.075, prior to coating and fixing with 4% formaldehyde on coverslips. Fixed cells were incubated with specific antibodies (α-FimA, 1:200 or α-FimP, 1:500) in 3% BSA for 1 h, followed by washing three times with PBS. Washed cells were then treated with AlexaFluor Plus 594 goat anti-rabbit IgG (Invitrogen) for 1 h, followed by washing three times with PBS. Coverslips were mounted on microscope slides with a drop of VectaShield mounting medium with DAPI (Vector Laboratories) and sealed with the nail polish before imaging by a fluorescence microscope (Keyence BZ-X810).

For quantification, more than 10 randomly selected areas of cells in each strain were used to determine the fluorescence levels of both channels red (AlexaFluor Plus 594) and blue (DAPI) using ImageJ. The results were expressed as averaged fluorescence levels of AlexaFluor, relative to DAPI signal, with statistical analysis performed with one-way ANOVA using GraphPad Prism; *, P<0.05; **, P<0.01; ***, P<0.001, and ns, P>0.05.

## DATA AVAILABILITY

All data are contained within this article. Atomic coordinates and structure factors for the SrtC2 structure were deposited into the Protein Data Bank as 8T28.

## Acknowledgments

We thank Dr. Robert Clubb (UCLA) and our lab members for their discussion and critical review of the manuscript.

## Author Contributions

C.C., HL. T.-T., J.O., and H.T.-T. conceptualization; C.C., HL. T.-T., and J.O. investigation; A.J. and H.T.-T. supervision; C.C. and H.T.-T. writing – original draft; C.C., HL. T.-T., J.O., A.D, and H.T.-T. writing – review and editing.

## Funding and Additional Information

Research reported in this publication was supported by the National Institute of Dental & Craniofacial Research (NIDCR) of the National Institutes of Health (NIH) under Award Number DE017382 (to H.T.-T) and in part by in part by federal funds from the National Institute of Allergy and Infectious Diseases, NIH, Department of Health and Human Services, under Contract 75N93022C00035 (to A.J). The use of SBC beamlines at the Advanced Photon Source is supported by the U.S. Department of Energy (DOE) Office of Science and operated for the DOE Office of Science by Argonne National Laboratory under Contract No. DE-AC02-06CH11357.

The content is solely the responsibility of the authors and does not necessarily represent the official views of the National Institutes of Health.

